# Pancreatic cancer-intrinsic HuR regulates the pro-tumorigenic properties of extracellular vesicles

**DOI:** 10.1101/2025.02.08.637077

**Authors:** Jennifer M. Finan, Yifei Guo, Alexandra Q. Bartlett, Matthew Reyer, Kevin Hawthorne, Margaret Haerr, Hen Halamish, Olayinka Lamikanra, Valerie Calvert, Canping Chen, Zheng Xia, Emanuel F. Petricoin, Rosalie C. Sears, Katelyn T. Byrne, Jonathan R. Brody

**Affiliations:** Department of Surgery, School of Medicine, Oregon Health & Science University, Portland, OR; Department of Cell, Developmental and Cancer Biology, Oregon Health & Science University, Portland, OR; Brenden-Colson Center for Pancreatic Care, School of Medicine, Oregon Health & Science University, Portland, OR; Knight Cancer Institute, Oregon Health & Science University, Portland, OR; Molecular and Medical Genetics, Oregon Health & Science University, Portland, OR; Applied Proteomics and Molecular Medicine, George Mason University, Manassas, VA; Biomedical Engineering Department, Oregon Health & Science University, Portland, OR

**Keywords:** Extracellular vesicles, pancreatic ductal adenocarcinoma, RNA-binding protein, human antigen R, endothelial cells

## Abstract

Pancreatic ductal adenocarcinoma (PDAC) tumors contain chaotic vasculature that limits immune surveillance and promotes early events in the metastatic cascade. However, current antiangiogenic therapies have failed in PDAC, and thus, it remains important to uncover mechanisms by which cancer cells signal to endothelial cells to increase angiogenesis. Our lab has shown that the tumor-intrinsic RNA-binding protein HuR (*ELAVL1*) plays an important role re-shaping the tumor microenvironment (TME) by regulating the stability and translation of cytokine encoding transcripts. Herein, we demonstrate that PDAC-intrinsic HuR influences endothelial cell function in the TME via extracellular vesicle (EV) signaling, an underexplored signaling axis in tumor progression. We found that HuR knockout (KO) tumors have impaired growth in an immunocompetent mouse model, and that administering purified wildtype (WT) EVs can increase tumor growth. Further, we observed that PDAC EVs contain HuR-dependent mRNA and protein cargoes relating to endothelial cell function and angiogenesis. Treatment of endothelial cells with HuR WT EVs strongly increased the expression of genes involved in barrier function and endothelial cell development, and directly increased their migratory and tube forming functions. In an immunocompetent orthotopic mouse model of PDAC, we showed that HuR increases endothelial cell presence and sprouting, while decreasing ICAM-1 expression. Importantly, we found utilizing a genetic EV reporter, that decreased ICAM-1 within WT tumors occurs in endothelial cells that have imported PDAC EVs, suggesting that this signaling axis is directly modulating endothelial cell behavior *in vivo*. Collectively, our data reveal a new role of HuR in EV signaling to endothelial cells, promoting angiogenesis while restricting endothelial cell leukocyte trafficking behavior.

## INTRODUCTION

Pancreatic ductal adenocarcinoma (PDAC) remains one of the most challenging solid tumors to diagnose and treat, in part, due to the complex and highly desmoplastic tumor microenvironment (TME)^1–3^. Embedded within the dense TME, endothelial cells comprise 2-10% of cells within primary PDAC tumors^4^. Endothelial cells are involved in angiogenesis, the formation of new blood vessels, leading to increased tumor vasculature. As this new vasculature is often abnormal, leading to leaky vessels that fuel tumor cell dissemination and metastasis, efforts have been made to inhibit angiogenic signaling pathways (e.g., VEGF, PDGF or FGF); however, clinical trials have been unsuccessful^5,6^. These findings highlight the need to strike a balance, inhibiting leaky vasculature while maintaining normal vasculature to ensure both immune surveillance and drug delivery^7^. Thus, it is important to define how PDAC cells signal to endothelial cells beyond classical growth factor signaling to receptor tyrosine kinases such as VEGFR, with the goal of identifying therapeutic intervention points to modulate endothelial cell function to improve patient outcomes.

One mechanism by which PDAC cells signal to surrounding cells is via extracellular vesicles (EVs), lipid bound particles that contain nucleic acids, metabolites, and proteins. Seminal work established the role of PDAC EVs in the metastatic cascade, setting up the pre-metastatic niche in the liver^8^. Importantly, it was illustrated that liver endothelial cells were among the highest importers of PDAC EVs, suggesting the PDAC EV-endothelial cell signaling axis plays a role in tumor progression^9^. Accordingly, recent work studying the fate of EVs utilizing genetic small EV reporters in mouse models found that endothelial cells import high levels of PDAC EVs *in vivo*^10,11^. Further, PDAC EV signaling to endothelial cells has been shown to increase endothelial cell proliferation and tube formation *in vitro*^12,13^. Despite these findings, little is understood about the tumor-intrinsic factors that are crucial for regulating PDAC EV cargoes, and thus their functional impact on recipient cells.

We previously demonstrated the RNA-binding protein, human antigen R (HuR, *ELAVL1*), directly impacts the TME composition, in part, through regulation of various cytokines and growth factors^14,15^. In fact, HuR regulates the production of VEGFA from PDAC cells, indicative of a possible intercellular signaling axis with endothelial cells^16^. We have shown that HuR post-transcriptionally regulates the translocation, stability, and translation of numerous key stress-response proteins such as PIM1, IDH1, and YAP1 within PDAC cells^15,17–19^. HuR contains three RNA recognition motifs that enable HuR to bind to a wide array of transcripts that contain AU-rich regions, enabling cells to rapidly respond to external stressors^20^. In PDAC and other solid tumor types, HuR translocation to the cytoplasm and increased levels of HuR target expression correlate with patient outcomes^14,15,21,22^. Further, HuR was shown to be within colon cancer EVs, supporting metastasis to the lung^23^. However, to date, no one has studied HuR’s impact on PDAC EV-mediated cell-cell signaling^24–26^. In the current study, we investigated how PDAC-intrinsic HuR impacts endothelial cells and vascular function via EV signaling. Here, we found PDAC EVs can increase tumor growth of HuR KO tumors. Using RNA-sequencing and quantitative proteomics we found that HuR regulates EV cargoes relating to endothelial cell biology, and that endothelial cell function is altered with WT vs. HuR KO EV treatment. Using an orthotopic mouse model of PDAC paired with a genetic EV reporter, PalmGRET, we found that endothelial cell abundance is dependent upon tumor-intrinsic HuR. Further, we found that EVs are imported by endothelial cells *in vivo*, independent of tumor HuR status, and that WT EV import leads to decreased ICAM-1 surface expression on endothelial cells. Together, these data define a role of PDAC-intrinsic HuR in regulating tumor EV cargoes that contribute towards dysfunctional vascular function *in vivo*.

## RESULTS

### Tumor-intrinsic HuR promotes tumor growth via EV signaling

Previous work from our group highlighted the role of tumor-intrinsic HuR regulating cell-cell communication through direct and indirect regulation of cytokines and growth factors^15,27^. Further, we found that *ELAVL1* deletion impairs tumor growth in an immunocompetent but not immunocompromised orthotopic mouse model of PDAC^15,27^. Based on previous work that tumor-intrinsic HuR regulates cell-cell signaling and existing literature highlighting the role of HuR in EV signaling in colorectal cancer, we assessed whether EVs derived from HuR proficient PDAC cells promote tumor growth^23,27^. As previously reported, we utilized CRISPR/Cas9 to genetically delete *ELAVL1* from PDAC cells derived from the Kras^G12D^;Trp53^R172H^;Pdx1-Cre (KPC) genetically engineered mouse model (**Fig. 1a**)^15^. We generated KPC mock, referred to as WT throughout, and HuR knockout (KO) single cell clones. We pooled three validated HuR KO clones for subsequent studies mixing 1:1:1. We found that HuR KO significantly reduces tumor growth in immunocompetent C57BL6 mice with orthotopic pancreatic tumors, validating our previous findings (**Fig. 1a-c**)^15^. To understand whether HuR plays a tumor-promoting role in EV signaling, we interrogated whether administering WT vs. HuR KO EVs to HuR KO tumor-bearing mice would alter the tumor phenotype (**Fig. 1g**).

**Figure 1:**
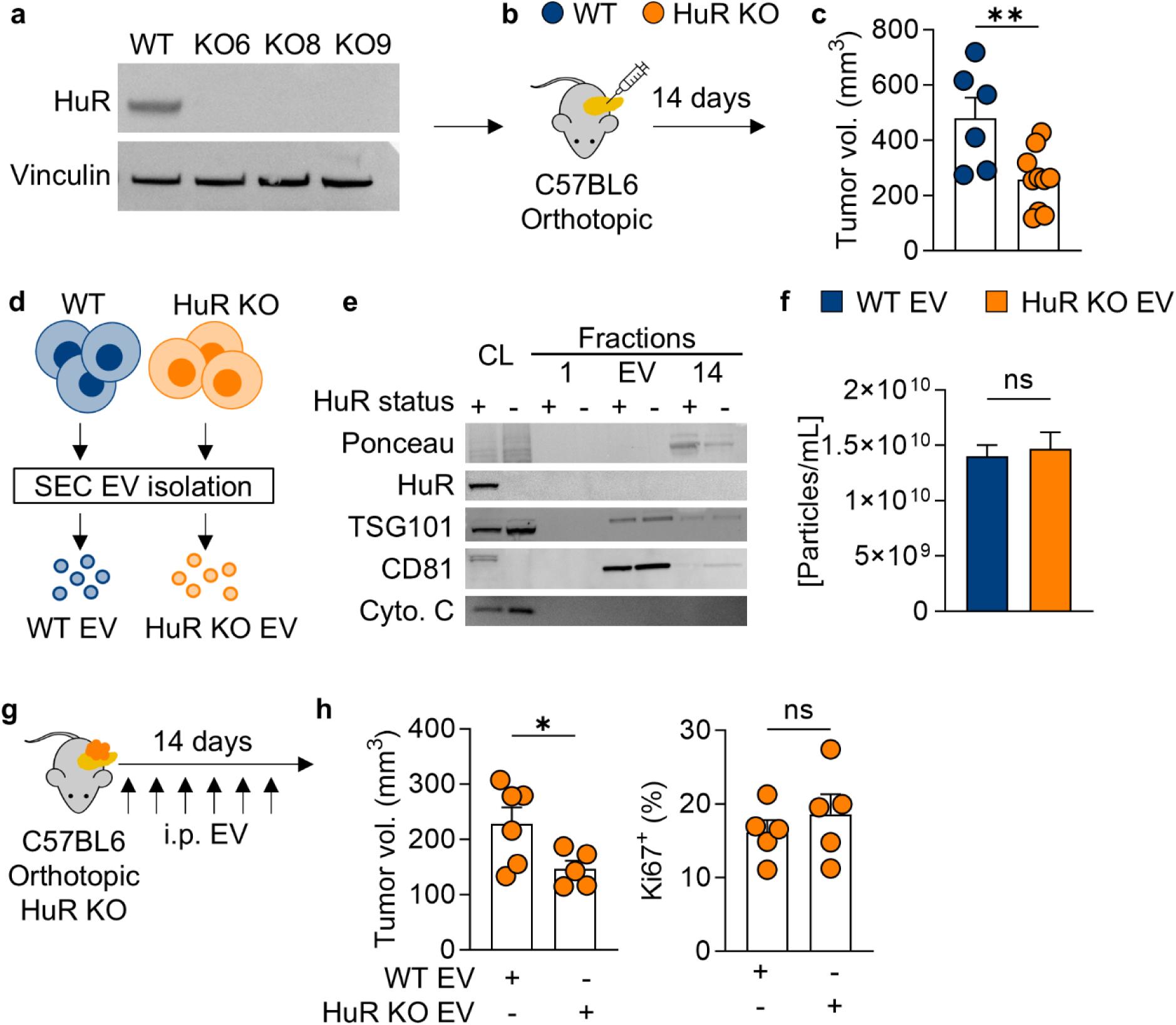
HuR WT EVs are tumor promoting. **(a)** Immunoblot validation of HuR expression in KPC WT and HuR KO clones 6-9 (loading control vinculin). **(b)** Schematic of pancreatic orthotopic implantation of KPC WT vs. HuR KO cells into C57BL6 mice and **(c)** tumor volume (mm^3^) after 14 days (WT *n* = 6, KO *n* = 10). **(d)** Schematic of EV isolation from KPC WT and HuR KO cells via size exclusion chromatography (SEC). **(e)** Immunoblot of cell lysates (CL) and SEC fractions from KPC WT (HuR status +) and HuR KO (HuR status -) cells. Blot probed for total protein (ponceau), HuR, EV markers (TSG101 and CD81) and cell lysate control (Cytochrome C). **(f)** Mean particle size concentration normalized to final cell number measurements via fluorescent nanoparticle tracking analysis. (*n* = 4). **(g)** Schematic of pancreatic orthotopic implantation of KPC KO cells into C57BL6 mice following i.p. injection of WT vs. HuR KO EVs every other day for 14 days and **(h)** tumor volume (mm^3^) after 14 days (WT *n* = 6, HuR KO *n* = 5) and immunofluorescence staining for proliferation via Ki67 staining (*n* = 5). *P* values were calculated using an unpaired two-tailed Student’s *t*-test. *, *P* < 0.05; **, *P* < 0.01; ***, *P* < 0.001; ns, not significant.

To isolate EVs from our mouse PDAC cell lines, we performed size exclusion chromatography (SEC) on 48-hour conditioned media collected from KPC WT or HuR KO cells following MISEV2023 guidelines (**Fig 1d-f**)^28^. EVs were confirmed to contain classical markers TSG101 and CD81, while being negative for HuR and cytochrome C, regardless of cell HuR status (**Fig. 1e**). We further validated our isolation utilizing fluorescent nanoparticle tracking analysis and found that KPC WT and HuR KO cells produce the same concentration of EVs (**Fig. 1f**). Next, we implanted KPC HuR KO cells into the pancreas of immunocompetent mice and administered WT or HuR KO EVs every other day for 14 days (**Fig. 1g**). We found that HuR KO tumors treated with WT EVs were significantly larger at the endpoint, suggesting that tumor-intrinsic HuR plays a tumor-promoting role via EV signaling (**Fig. 1h**). Further, we found that WT EV and HuR KO EV treated tumors had equivalent Ki67 positivity by immunofluorescence staining, suggesting that the differences in tumor size are not due to changes in proliferation, but due to alterations in the TME. Importantly, we found that HuR is not present within WT EVs, indicating that HuR is having a tumor-promoting effect likely though cargoes that are different in EVs derived from WT vs. HuR KO cells (**Fig. 1e**).

### HuR impacts PDAC EV cargoes relating to endothelial cell functions

To interrogate how HuR can promote tumor growth via EV signaling, we next sought to characterize the cargoes of EVs from WT vs. KO PDAC cells. We utilized the human PANC-1 PDAC cell line (KRAS^G12D^, TP53^R273H^) to provide human disease relevance, and contains a similar mutational profile as the KPC-8069 cell line (KRAS^G12D^, TP53^R172H^). We generated PANC-1 mock and HuR knockout (KO) single cell clones and validated their loss of HuR at the protein level (Fig. S1). For subsequent studies pooled PANC-1 mock, referred to throughout as WT, and PANC-1 HuR KO clones 1-6 were pooled at equal ratios. We isolated EVs from conditioned media collected from PANC-1 WT or HuR KO cells utilizing size exclusion chromatography following MISEV2023 guidelines (**Fig. 2a**)^28^. Next, we validated our EV isolation utilizing protein content, showing a first protein peak in fractions 7-10 that contains EVs followed by a larger peak in fractions 12-14, indicative of increased albumin proteins (Fig. S1). EVs were confirmed to contain classical markers TSG101 and CD81, while being negative for HuR and cytochrome C, regardless of cell HuR status (**Fig. 2b**).

**Figure 2:**
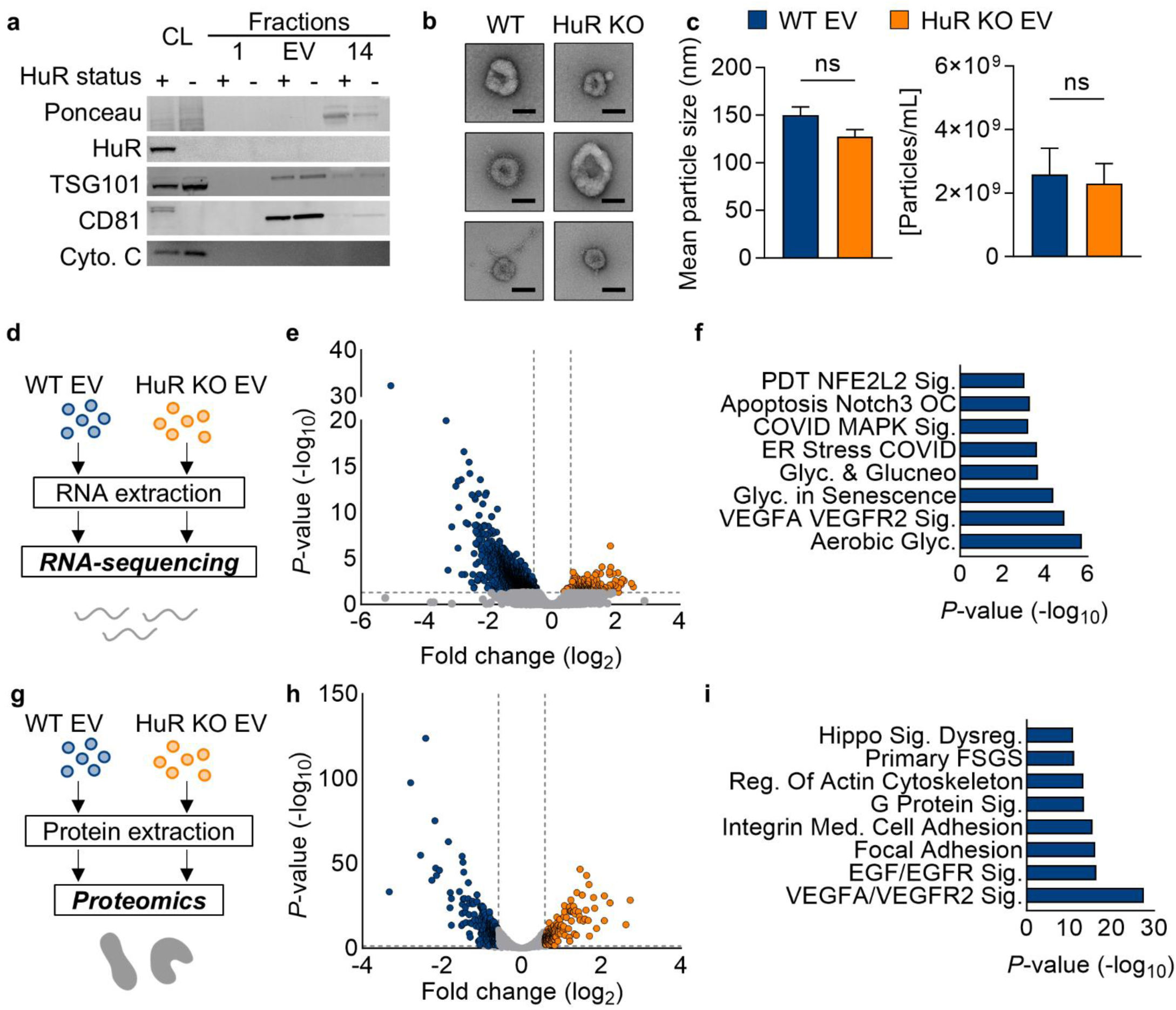
HuR impacts PDAC EV cargoes relating to endothelial cell functions. **(a)** Immunoblot of cell lysates (CL) and SEC fractions from PANC-1 WT (HuR status +) and HuR KO (HuR status -) cells. Blot probed for total protein (ponceau), HuR, EV markers (TSG101 and CD81) and cell lysate control (Cytochrome C). **(b)** Electron microscopy validation of EV isolation (scale bar = 50 nm). **(c)** Mean particle size and concentration normalized to final cell number measurements via fluorescent nanoparticle tracking analysis. *P* values were calculated using an unpaired two-tailed Student’s *t*-test (*n* = 4). **(d)** Schematic of RNA-sequencing of WT vs. HuR KO EVs. **(e)** Volcano plot of differentially expressed genes in WT (blue, left) vs. HuR KO (orange, right) EVs (*n* = 4). **(f)** The top 8 enriched pathways in HuR WT EVs. **(g)** Schematic of isobaric-labeling quantitative proteomics of WT (blue, left) vs. HuR KO (orange, right) EVs (*n* = 3). **(h)** Volcano plot of differentially abundant proteins in HuR WT vs. KO EVs (*n* = 3). **(i)** The top 8 enriched pathways in HuR WT EVs.

We further validated our EV isolation via electron microscopy and fluorescent nanoparticle tracking analysis (**Fig. 1c**, Fig. S1). We found no difference in the size or secretion of the particles isolated by size exclusion chromatography (**Fig. 2c**). Next, we assessed whether there were different cargoes within these EVs. Upon mRNA-sequencing of PANC-1 WT vs. HuR KO EVs, we found that tumor-intrinsic HuR impacts the mRNA cargoes within PDAC EVs (**Fig. 2d**, Supplementary Data 1). Specifically, PANC-1 WT EVs contained HuR-dependent mRNAs within pathways relating to cellular metabolism and endothelial cell biology (**Fig. 2e-f**). Given that HuR impacts mRNA cargoes within EVs, we performed isobaric-labeling quantitative proteomics analysis on these EVs to look for changes in protein cargoes. We again found a significant number of HuR-dependent proteins (**Fig. 2g-i**, Supplementary Data 2). Upon pathway analysis we found that pathways implicating endothelial cell function were significantly altered, consistent with changes in mRNA cargo. We additionally performed a reverse phase protein array (RPPA) to better interrogate signaling pathways regulated in a tumor-intrinsic HuR-dependent manner in these EVs. We found that the most differentially abundant proteins and phospho-proteins in our EVs were again related to endothelial cell biology (Fig. S1). Together, these data confirm that tumor-intrinsic HuR regulates EV cargoes that may regulate endothelial cell function.

To determine whether HuR is important in endothelial cell function within the TME in human patients with PDAC, we correlated *ELAVL1* expression to the presence of endothelial cells from a publicly available single cell RNA-sequencing dataset^29^. Strikingly, *ELAVL1* expression correlated with the abundance of endothelial cells in these data, and there were a significantly higher number of endothelial cells in the top quartile of *ELAVL1* expressing patients comparted to the bottom quartile (**Fig. 3a-b**). These data suggest that PDAC cell-intrinsic HuR may be regulating endothelial cell presence within PDAC tumors, in part via EV signaling.

**Figure 3:**
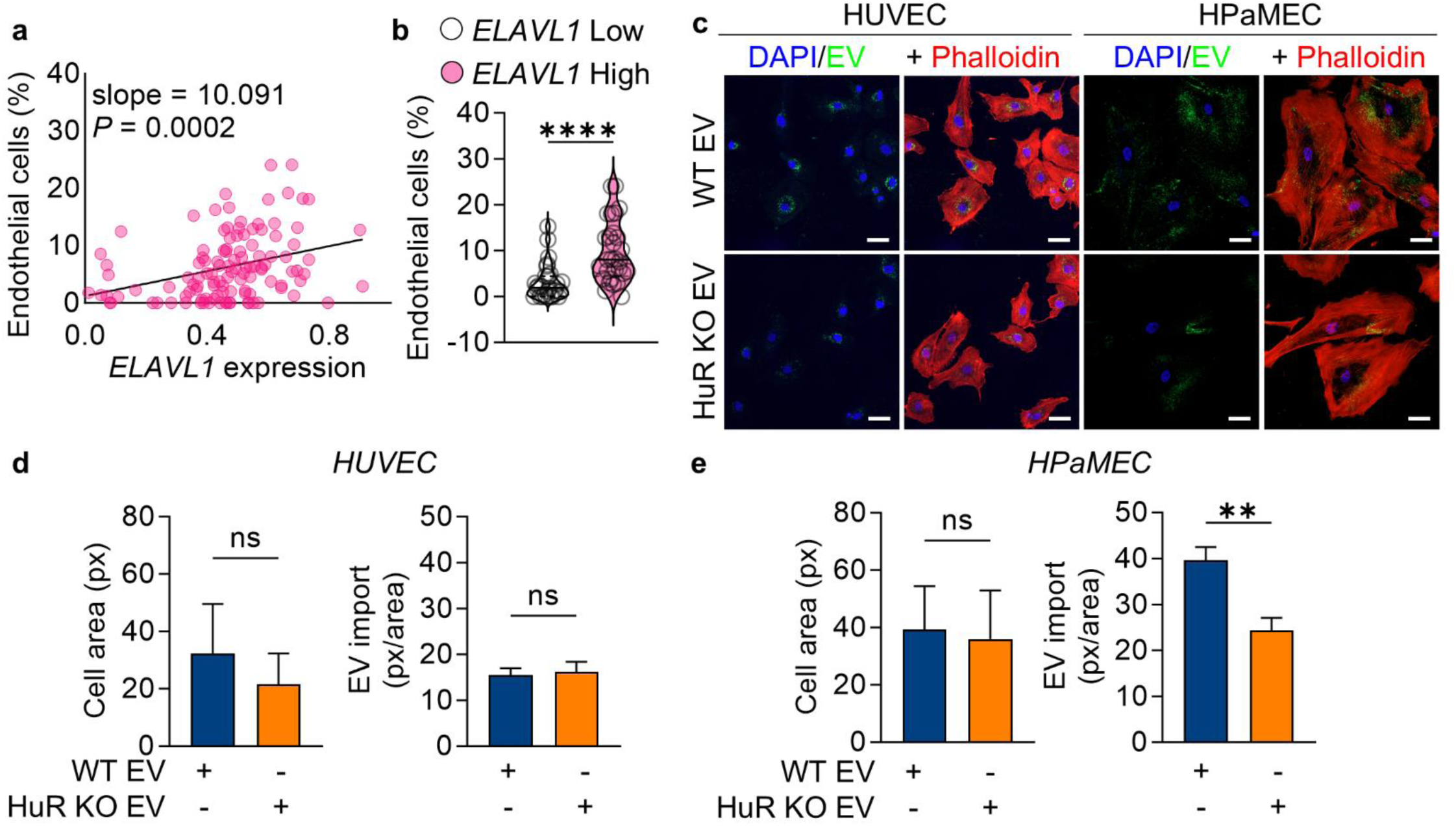
Endothelial cell abundance correlates with *ELAVL1* expression and endothelial cells import PDAC EVs. **(a)** Correlation between *ELAVL1* expression in PDAC cells and presence of endothelial cells in publicly available patient single cell RNA-sequencing. **(b)** Within this cohort, comparison of endothelial cell percentage in patients with the highest and lowest quartile of *ELAVL1* expression. **(c)** Representative images of HUVECs and HPaMECs treated with PKH67 labeled PANC-1 WT vs. HuR KO EVs for 4 hours with stained nuclei (blue, DAPI) and cytoskeleton (phalloidin, red) (scale bar = 50 µm). **(d)** Relative cell area of HUVECs and **(e)** HPaMECs after 4-hour EV treatment and levels of EV internalization (*n* = 3).

### PDAC EVs are readily imported by endothelial cells in vitro

To assess whether endothelial cells import human PDAC EVs, we optimized *in vitro* EV import studies leveraging the lipophilic dye, PKH67^8,30,31^. We treated endothelial cells with PKH67 alone or PKH67 labeled EVs over 8 hours and detected that cells import a significant level of EVs by 4 hours, after which the lipid signal becomes diffuse within the cell as the lipids are recycled (Fig. S2a). Further, we found that EV import is concentration dependent, thus for subsequent EV import studies, we treated cells with the concentration of EVs we found PDAC cells to have in the media after 48 hours, roughly 300 EVs/cell (Fig. S2b). Finally, we validated that our technique monitors EV uptake rather than lipid aggregates as either 4°C or dye only treated conditions have no EV signal after 4 hours (Fig. S2c). Using this approach, we observed that both human umbilical vein endothelial cells (HUVECs) and human pancreatic microvascular endothelial cells (HPaMECs) import PDAC EVs (**Fig. 3c-e**). Interestingly, we found that HPaMECs, but not HUVECs, imported significantly fewer HuR KO EVs than WT EVs. These results are concordant with reports that EV import may be a regulated process by which tissue or cell specific receptors facilitate EV import through receptor-ligand binding^32–34^. Together these data validate previous findings that endothelial cells indeed import PDAC EVs *in vitro*^10,12,13^.

### HuR WT EVs directly alter endothelial cell function in vitro

Next, we sought to determine whether WT vs. HuR KO EVs differentially impact endothelial cell function. We treated HPaMECs with PANC-1 WT vs. HuR KO EVs for 24 hours and performed RNA-sequencing on the HPaMECs (**Fig. 4a**). We found that transcripts in pathways relating to barrier function were significantly altered in the WT vs. KO EV treated endothelial cells (**Fig. 4b-c**, Supplementary Data 3). Next, to assess whether these mRNA changes in EV treated endothelial cells resulted in functional changes, we treated HPaMECs or HUVECs for 24 hours in culture prior to measuring endothelial cell migration, tube formation, and barrier function (**Fig. 4d-f**, Fig. S3). We observed that HPaMECs and HUVECs migrate more when treated with WT EVs compared to HuR KO EVs, and that these EV treatments differentially impacted barrier function and tube formation. Specifically, we found that WT EV treated endothelial cells have improved barrier function, which can be either tumor-promoting due to decreased immune infiltration or tumor-suppressing due to decreased tumor cell intravasation (**Fig. 4e**, Fig. S3b)^35^. Further, we found that HuR KO EV treated endothelial cells form fewer tubes than WT EV treated endothelial cells as indicated by the decrease in total tube length, number of branching points, and number of loops after 8 hours (**Fig. 4f**, Fig. S3c-d). These observations validate the ability of PDAC HuR to impact endothelial cells via EV signaling in a human *in vitro* model.

**Figure 4:**
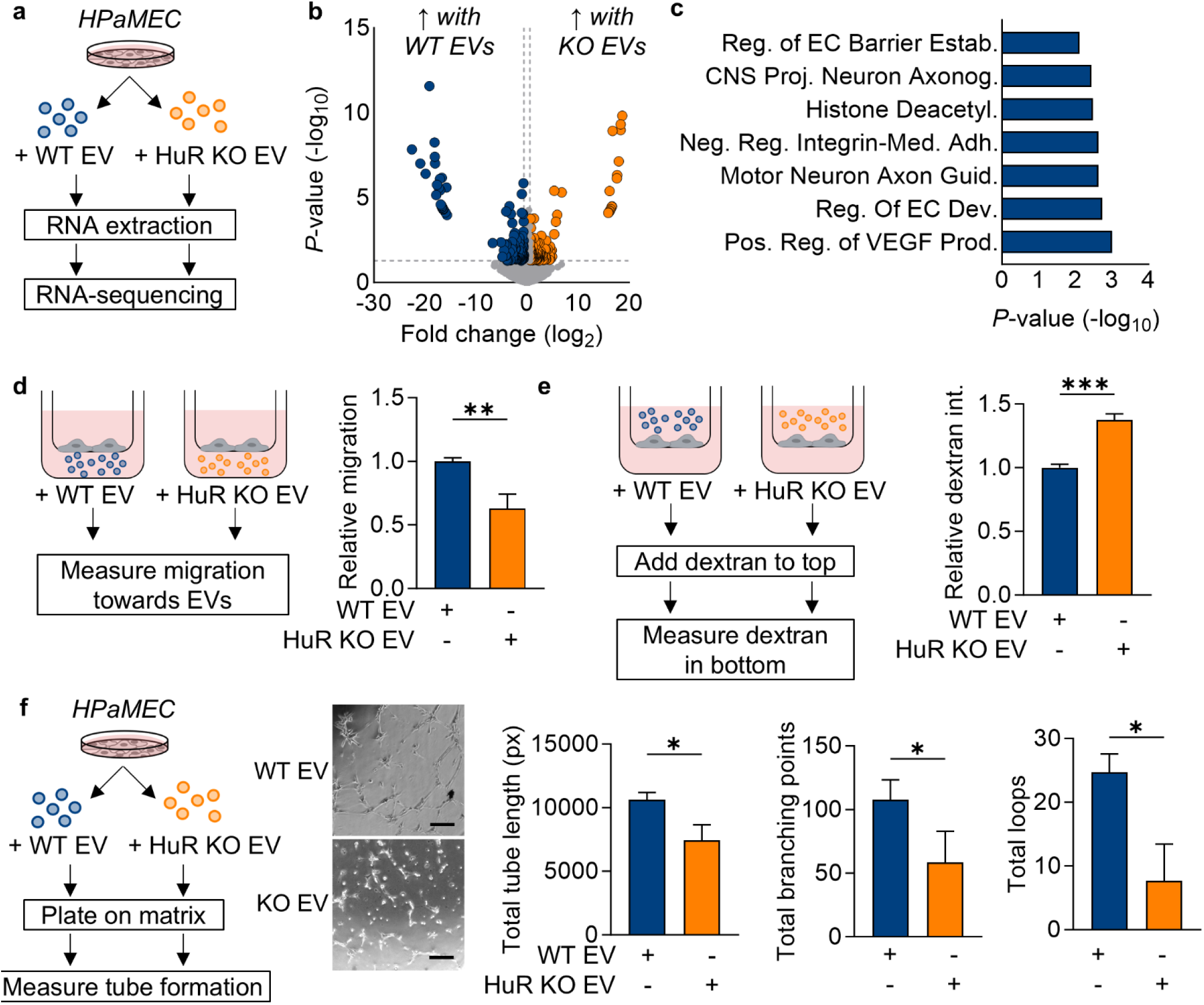
WT EVs directly impact endothelial cell transcriptome, migration, barrier function, and tube formation. **(a)** Schematic of RNA-sequencing HPaMECs treated with WT vs. HuR KO EVs for 24 hours. **(b)** Volcano plot of differentially expressed genes in HPaMECs treated with PANC-1 HuR WT vs. KO EVs for 24 hours (*n* = 3). **(c)** Gene Ontology biological process analysis of the top 7 pathways enriched in WT EV treated HPaMECs (*n* = 3). **(d)** Transwell migration of HPaMECs towards PANC-1 WT vs. HuR KO EVs over 24 hours and quantified with crystal violet staining (*n* = 3). Functional analysis of HPaMECs treated with media alone, PANC-1 WT or HuR KO EVs for 24 hours and monitored for **(e)** monolayer permeability quantified by dextran movement across the endothelial cell monolayer and **(f)** tube formation quantified for total tube length (px), total branching points and total loops (*n* = 3) (scale bar = 250 µm). *P* values were calculated using an unpaired two-tailed Student’s *t*-test. *, *P* < 0.05; **, *P* < 0.01; ***, *P* < 0.001; ns, not significant.

### PDAC-intrinsic HuR is necessary for endothelial cell recruitment

Based on our findings that PDAC-intrinsic HuR correlates with endothelial cells abundance in patient PDAC samples, we aimed to determine whether our KPC mouse model could recapitulate this. We implanted immunocompetent C57BL6 mice with subcutaneous WT vs. HuR KO KPC cells and allowed tumors to develop over 30 days (Fig. S4). Based on our findings that HuR WT EVs can modulate endothelial cell function *in vitro*, we assessed the presence of endothelial cells in these mouse tumors. We found that tumor cell loss of HuR significantly decreased the presence of endothelial cells as stained for by endomucin (Fig. S4a).

Next, we validated these findings in an orthotopic pancreatic mouse model by implanting WT and HuR KO cells into the pancreas of C57BL6 mice and allowing tumors to develop over 14 days (**Fig. 5a**). Similarly to the subcutaneous setting, orthotopic HuR KO tumors had 30% fewer endothelial cells as quantified by immunofluorescence. Importantly, the function in addition to presence of vasculature is crucial in impacting tumorigenesis, thus we aimed to assess whether the vasculature in WT vs. HuR KO tumors functioned similarly. We performed ultrasound imaging on WT vs. HuR KO tumor-bearing mice at 13 days after implantation using power doppler 3D mode to measure percent vascularity, indicative of the relative vascular density within tumors. In accordance with decreased endothelial abundance via immunofluorescence staining, we found that there was a decrease in percent vascularity in HuR KO tumors compared to WT (**Fig. 5b**). Next, we retro-orbitally injected fluorescently labeled lectin to label functional vasculature within tumors and assessed percent functional vasculature by staining for endomucin. We observed that both WT and HuR KO tumors had approximately 50% functional perfused vasculature (Fig. S4b).

**Figure 5:**
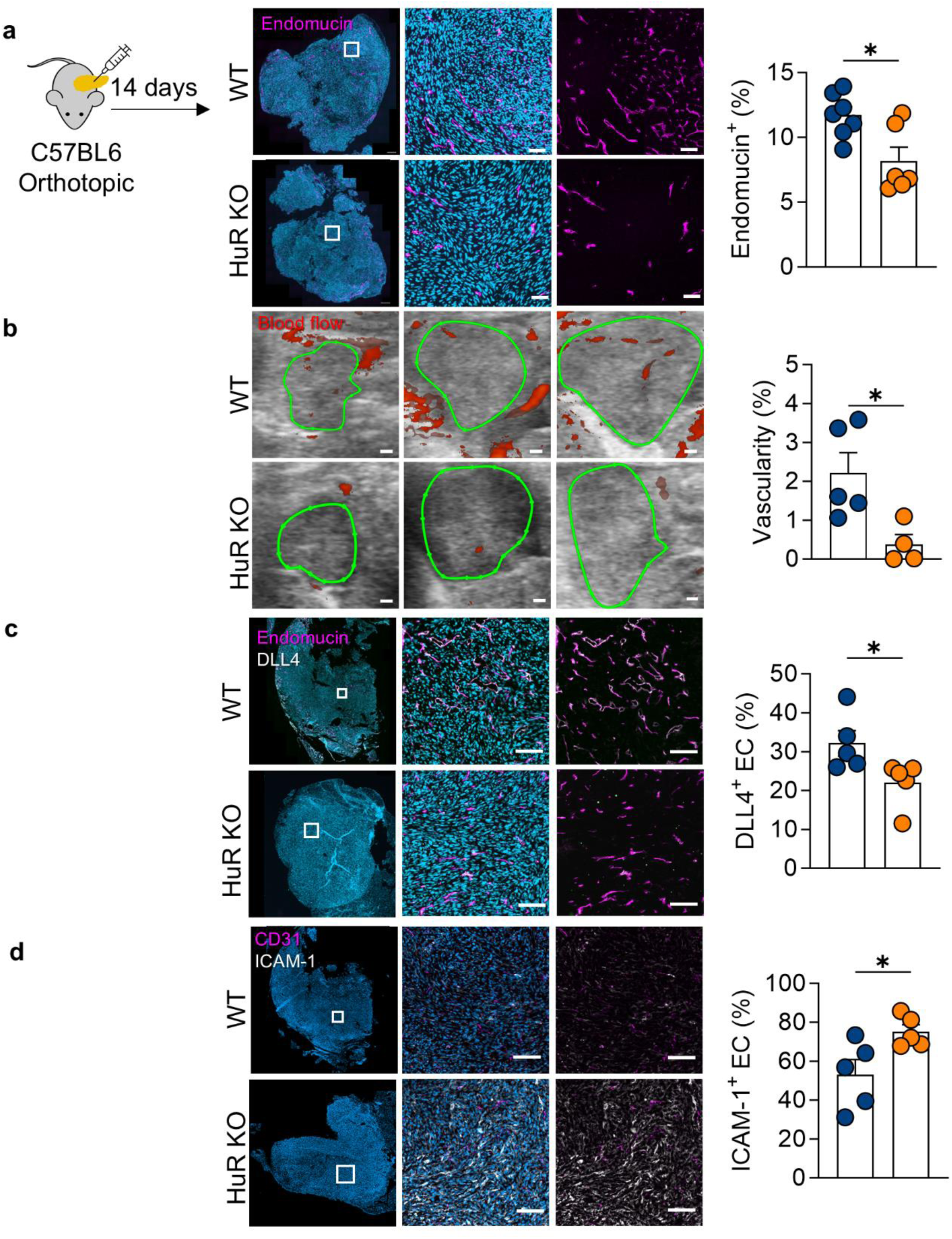
Tumor-intrinsic HuR is promotes endothelial cell recruitment and sprouting in PDAC tumors. **(a)** KPC HuR KO cells implanted orthotopically into the pancreas (*n* = 7) of immunocompetent C57BL6 mice decreased endothelial cell presence compared to WT tumors as quantified by staining for endomcucin (magenta) and nuclei (DAPI, teal) (scale bar = 100 µm). **(b)** Ultrasound power doppler imaging of orthotopic KPC WT and HuR KO tumors to quantify percent vascularity at 13 days post implantation (*n* = 4) (scale bar = 1 mm). **(c)** Immunofluorescence of tumors co-staining for endothelial cells (endomucin, magenta) and sprouting (DLL4, white) (scale bar = 100 µm). **(d)** Immunofluorescence of tumors co-staining for endothelial cells (CD31, magenta) and ICAM-1 (white) (scale bar = 100 µm). *P* values were calculated using an unpaired two-tailed Student’s *t*-test. *, *P* < 0.05; **, *P* < 0.01; ***, *P* < 0.001; ns, not significant.

To further interrogate the function of the vasculature within these tumors, we stained for a marker of endothelial cell sprouting, DLL4. We found that there are 45% more sprouting endothelial cells in WT tumors than HuR KO tumors, indicative of increased vascular remodeling (**Fig. 5c**). Next, we evaluated the surface expression of ICAM-1, an important cell surface glycoprotein that serves as a scaffold for leukocytes binding, aiding trans-endothelial cell migration^36,37^. We observed that ICAM-1 surface expression on endothelial cells increased in HuR KO tumors, suggesting an increase in inflammatory activation in HuR KO tumor endothelial cells (**Fig. 5d**). Taken together, these data suggest that PDAC-intrinsic HuR increases endothelial cell abundance and sprouting, while decreasing the ability of the vasculature to aid leukocyte trafficking within the TME.

### PDAC HuR impacts endothelial cells in vivo directly via EV import

EV signaling dynamics are impacted by many physiological factors including matrix stiffness and pH, which are greatly different between *in vitro* culture and PDAC tumors^38–41^. To study the role of PDAC-intrinsic HuR in EV signaling in a physiological setting, we employed an established genetic EV reporter, PalmGRET, which incorporates GFP-nLuc into the inner bilayer of cellular lipid bilayers via a palmitoylation sequence (**Fig. 6a**)^42,43^. Leveraging this model, we could determine whether lipid bound particles, including EVs, derived from WT and HuR KO PDAC cells are trafficked to endothelial cells within the TME. Importantly, this model relies on endogenous production of EVs from tumor cells within a physiological setting with tumor-related stressors, rather than administering exogenously produced EVs. We validated that this reporter in KPC cells leads to the production of GFP^+^ and nLuc^+^ EVs utilizing two independent techniques (**Fig. 6b**, Fig. S5a). Further, addition of this construct did not alter the *in vitro* growth of these cells, of note, *in vivo* tumor size was altered yet not significantly (**Fig. 6c**, Fig. S5b). This may be due to the immunogenicity of exogenous proteins such as GFP and nLuc, a caveat to utilizing any currently available genetic EV reporter^44^. We generated WT and HuR KO cells expressing the PalmGRET reporter, and FACS sorted them to ensure they have equivalent GFP signal (Fig. S5c), as confirmed in each subsequent experiment via immunoblotting for nLuc (**Fig. 6d**). Next, these cells were utilized to assess which stromal cells import PDAC WT vs. HuR KO EVs *in vivo* by establishing orthotopic pancreatic tumors in immunocompetent mice (**Fig. 6e**).

**Figure 6:**
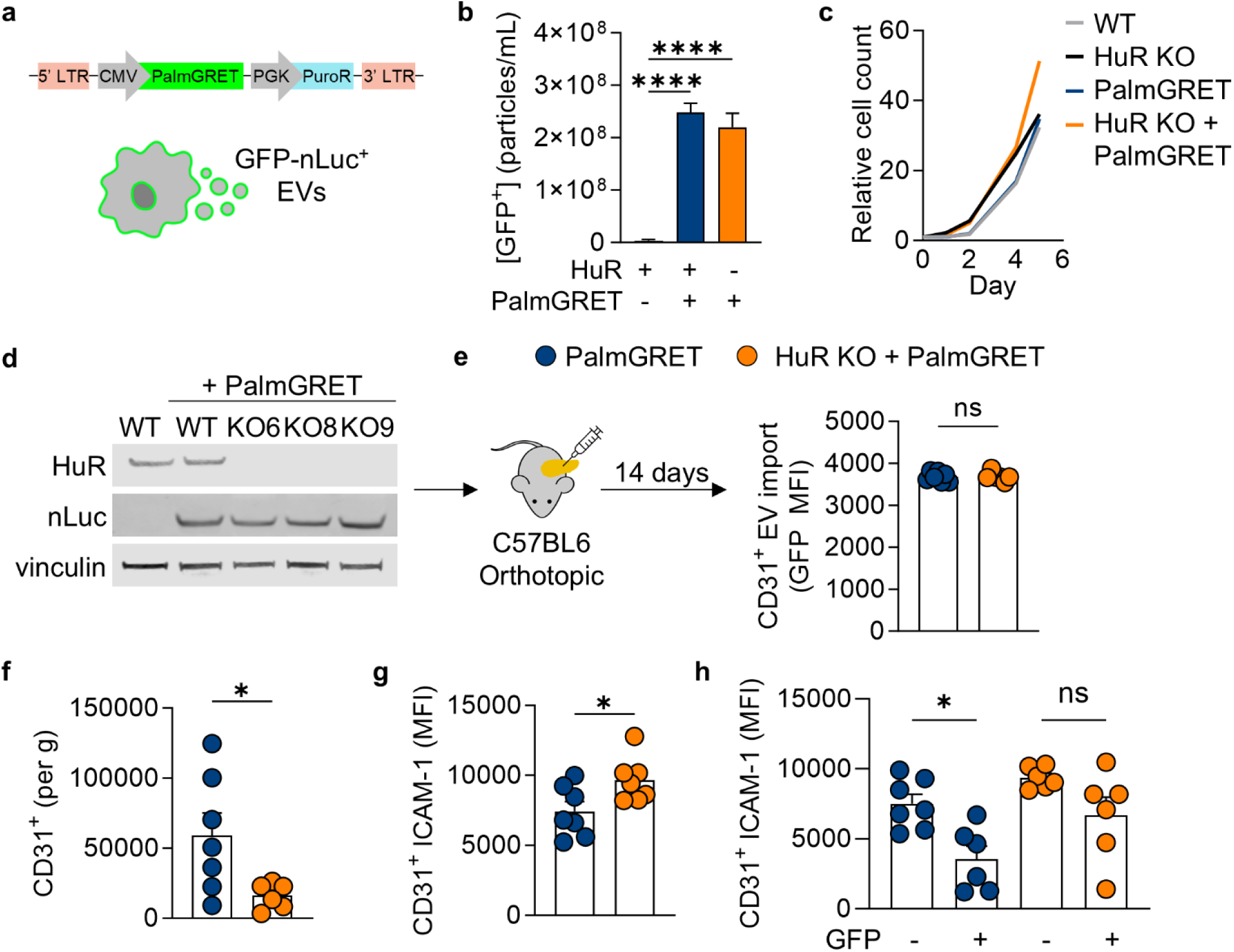
Tumor-intrinsic HuR alters vascular and lymphatic endothelial cell function. **(a)** Schematic of the PalmGRET reporter labeling all inner-leaflets of membranes with GFP-nLuc via integration of a palmitoylation sequence. The construct for the PalmGRET EV reporter labeling the inner leaflet of all cellular membranes and secreted particles. **(b)** PalmGRET expressing cells produce GFP^+^ EVs detected by fluorescent nanoparticle tracking analysis (*n* = 3). **(c)** Cell growth of PalmGRET expressing KPC cells over 6 days. **(d)** Immunoblot of cell lysates utilized for mouse studies probed for HuR, nLuc and loading control vinculin. **(e)** PalmGRET expressing cells were orthotopically implanted into C57BL6 mice and tumor were harvested at 14 days for flow cytometry. Levels of EV import (GFP MFI) in endothelial cells (CD45^-^CD31^+^) were equal in WT and HuR KO tumors *(n* = 7)*. **(e)*** Total endothelial cell (CD45^-^CD31^+^) presence as quantified by cells per gram of tumor in WT vs. HuR KO tumors. **(g)** ICAM-1 geometric mean surface expression of ICAM-1 in endothelial cells (CD45^-^CD31^+^) of WT vs. HuR KO tumors (*n* = 7). **(h)** ICAM-1 geometric mean surface expression (MFI) of ICAM-1 in endothelial cells of WT vs. HuR KO tumors that have (GFP^+^) or have not (GFP^-^) imported PDAC EVs. *P* values were calculated using an unpaired two-tailed Student’s *t*-test or an ordinary one-way ANOVA (panel h only). *, *P* < 0.05; **, *P* < 0.01; ***, *P* < 0.001; ns, not significant.

We sought to determine the levels of PDAC EV import by endothelial cells within the TME. To address this, we performed flow cytometry on tumors, staining for stromal cell markers and evaluating levels of GFP (Supplemental Table 1 & 2). GFP geometric mean fluorescence intensity (MFI) within endothelial cells (CD45^-^ CD31^+^) in WT and HuR KO tumors were similar, indicating that endothelial cells within both tumors are importing PDAC EVs, irrespective of PDAC cell-intrinsic HuR (**Fig. 6e**). In concordance with prior studies, we observed that other cells, including cancer associated fibroblasts and dendritic cells within the TME import PDAC EVs in both WT and HuR KO tumors (Fig. S5d). Importantly the high GFP signal in macrophages may be indicative of both EV import and phagocytosis, as these two events cannot be distinguished based upon GFP signal alone. Our data aligned with previously published studies, where cancer associated fibroblasts and dendritic cells are major importers of PDAC EVs^11^. Together, these data suggest that the tumor-promoting role of HuR via EV signaling occurs due to the impact of the EV cargoes on endothelial cells rather than rates of EV import.

In addition to EV import rates, we calculated the number of endothelial cells per gram of tumor and found that in accordance with previous studies, HuR KO tumors have significantly fewer endothelial cells (CD45^-^CD31^+^) (**Fig. 6f**). Further, we observed that ICAM-1 surface expression on endothelial cells was lower in WT tumors than in HuR KO tumors, in concordance with our findings in the KPC model without PalmGRET (**Fig. 6g**). Importantly, we found that within the WT tumors, endothelial cells that had imported a PDAC EV (GFP^+^) had lower ICAM-1 surface expression when compared to endothelial cells from WT tumors that had not imported a PDAC EV (GFP^-^) (**Fig. 6h**). However, the endothelial cells in HuR KO tumors had elevated ICAM-1 surface expression, irrespective of whether they were GFP^+^. These data suggest that HuR-dependent cargoes within WT EVs lead to decreased ICAM-1 surface expression on endothelial cells. These findings highlight an important role of HuR in regulating the function of the vasculature within the TME, which can impact tumor progression, immune surveillance, and response to therapy^45,46^.

## DISCUSSION

Intercellular communication has long been understood to play a role in many facets of tumor progression, enabling genetically mutated cancer cells to co-opt normal tissues^1,2,47^. More recently, EV signaling has become appreciated as a coordinated signaling mechanism within and across tissues with the improvement of genetic EV reporters to study this signaling axis *in vivo*^11,42,48,49^. In PDAC, it has been shown that EVs are crucial in establishing a pre-metastatic niche and have the potential to play a role early in the metastatic cascade via signaling to endothelial cells and fibroblasts within the TME^8,9,11^. However, little has been done to understand the key cargoes and regulators of these cargoes in PDAC EV signaling. Our group and others have established the role of the aberrantly regulated RNA-binding protein HuR in numerous cell-intrinsic stress-adaptive processes and, more recently, in cell-cell signaling within TME^27^. Herein, we aimed to determine the role of PDAC-intrinsic HuR in EV signaling to the TME and found that PDAC EVs are tumor-promoting in an HuR dependent manner, likely due to its impact of endothelial cell function.

We found that HuR KO cells produce EVs at the same rates and size; however, there are significant changes to the mRNA and protein cargoes, both of which can induce functional changes in recipient cells. Of note, we found that HuR itself is not present within PDAC EVs, rather HuR regulates cargoes that are within EVs. We found that both mRNAs and proteins relating to metabolism and endothelial cell function were the most impacted. We further found that EV biogenesis proteins are among the proteins unchanged with HuR status, supporting our findings that HuR does not change EV secretion rates. We and others have shown that PDAC EVs are imported by endothelial cells *in vitro* and *in vivo*. Our findings corroborate work performed in two other independent genetic EV tracking models that EVs are imported by specific cell types, rather that equally across all cells in the TME^11^. However, by leveraging the PalmGRET model we were able to track the broad heterogenous population of EVs, rather than a subset of lipid-bound particles^48^. Although there are changes to the mRNA and protein cargoes with the loss of tumor-intrinsic HuR, there are no changes to *in vivo* PDAC EV import. This suggests that the molecules involved in PDAC EV import are HuR-independent, whereas the function of these EVs on endothelial cells is HuR-dependent, as observed in both RNA-sequencing and phenotyping studies on HPaMECs with EV treatment.

Using murine PDAC models, we demonstrated that PDAC HuR is necessary for endothelial cell recruitment within the TME. These data are concordant with the correlation between *ELAVL1* expression and endothelial cell abundance in patient samples. Importantly we know that vascular function rather than presence correlates with patient outcomes^7^. Endothelial cells are dynamic and crucial cells making up the barriers of vasculature throughout the body. As widely reported tumor vasculature is often dysfunctional, further exacerbating hypoxia within tumors and promoting more chaotic angiogenesis^50^. In both our WT and HuR KO tumors we found that only 50% of the vasculature was functional, further, within WT tumors there was an increase in DLL4 expressing tip cells. The high level of DLL4 expressing cells is another indicator that the vasculature is chaotic and uncontrolled^51^. The implications of this chaotic vasculature may promote metastasis and/or immune evasion in PDAC tumors^7^.

There is an important interplay between vasculature and immune cells that allows for proper immune surveillance within tissues^45,46^. During inflammatory responses, endothelial cells express molecules to promote leukocyte trans-endothelial cell migration such as ICAM-1 and reduce their barrier permeability^52^. Our data suggest that in WT tumors there is impaired ICAM-1 surface expression when compared to HuR KO tumors. Importantly, it is specifically the endothelial cells that have imported PDAC EVs in WT tumors that have a decrease in ICAM-1 surface expression compared to endothelial cells within the same tumor that have not imported an EV. The decrease in ICAM-1 surface expression could be associated HuR-dependent EV cargoes that suppress Wnt or NF-κB signaling, both of which increase ICAM-1 expression in endothelial cells^53,54^. Specifically, protein cargoes SOD3, SFRP1, DKK1 were significantly enriched in WT EVs and have been implicated in suppression of Wnt or NF-κB signaling^55–59^. Together, these data implicate PDAC EV signaling in regulating endothelial cells ICAM-1 surface expression, which has potential implications for immune trafficking. Future studies are warranted to understand whether the vascular remodeling can directly regulate immune function within our model.

To conclude, our findings define HuR as an important regulator of tumor derived-EV cargoes, specifically within the PDAC EV-endothelial cells signaling axis. We showed that WT EVs are tumor-promoting and can directly increase endothelial cell migration and tube formation. Moreover, using *in vivo* modeling we show that EVs are imported by endothelial cells and have distinct functional changes within WT tumors. The direct mechanism of how PDAC EVs induce these phenotypes in endothelial cells is most likely due to many pathways orchestrated by HuR. Overall, our characterization of HuR-dependent EV cargoes and the vasculature remodeling within WT tumors contributes to a deeper understanding of the PDAC EV signaling axis and how tumor-intrinsic HuR directly effects the TME. This work lays the groundwork for identifying future vascular targets within the PDAC TME to improve the delivery and efficacy of current therapies targeting PDAC.

## MATERIALS AND METHODS

### Cell lines

Human PDAC cell lines (PANC-1, MIA PaCa-2) were obtained from the American Type Culture Collection (ATCC) and cultured in DMEM supplemented with 10% fetal bovine serum (FBS) and 1% penicillin-streptomycin. Cells were authenticated via short tandem repeat analysis at the OHSU Gene Profiling Shared Resource and routinely tested for mycoplasma. Mouse PDAC cell line (KPC-8069) was gifted by Dr. Tony Hollingsworth (University of Nebraska Medical Center) and cultured in DMEM. Endothelial cells used were human pancreatic microvascular endothelial cells (HPaMECs) purchased from ScienCell Research Laboratories and human umbilical vein endothelial cells (HUVECs) purchased from ATCC. Endothelial cells were cultured in endothelial cell medium (ScienCell #1001) in flasks coated with fibronectin (2 µg/cm^2^) or Quick Coating Solution (Angio Proteomie #CAP-01). All cell lines were cultured at 5% CO_2_ at 37°C in a humidified atmosphere.

### Extracellular vesicle isolation

Cells were plated at 2-3.5×10^6^ per 15 cm dish in 20 mL of DMEM supplemented with 10% FBS and 1% penicillin-streptomycin. The next day, media was changed to DMEM supplemented with 10% EV-depleted FBS. After 48 hours, conditioned media was collected and spun at 2000 x g for 10 minutes to remove cell debris. Next, the media was concentrated utilizing Amicon Ultra-15 Centrifugal Filer Unit (Millipore #UFC9100) filter columns down to 1 mL. Size exclusion chromatography (SEC) was performed utilizing Izon qEV1 columns with 0.22 µm filtered PBS, and EVs were collected in pooled fractions 7-10. Simultaneously, cells are trypsinized following conditioned media collection for cell counting and pelleting for EV particle concentration normalization and controls via immunoblotting.

### Fluorescent nanoparticle tracking analysis

Lipid dye, Di-8-ANEPPS (Biotium #61012), was prepared diluting 1:100 in PBS with 0.05% Pluronic F-127. 20 µL of each sample was incubated with 1 µl of dye for 15 minutes and immediately diluted with 979 µl H_2_O. Samples were run on the ZetaView utilizing the 488 nm laser with the 500 nm filter at 11 positions at 25°C. Particle concentration was reported and normalized to the cell count of donor cells.

### Transmission electron microscopy

SEC fractions were submitted to the OHSU Multiscale Microscopy Core for sample processing and imaging. Samples were placed on 200 mesh grids coated with carbon and formvar for 3 minutes. The grids were rinsed three times in water and exposed to 1% (w/v) uranyl acetate for 3 minutes. The grids were blotted dry and imaged on a ThermoFisher Scientific Tecnai TEM transmission electron microscope operated at 120kV equipped with an AMT Nanosprint12 camera.

### Immunoblotting

Cell pellets were washed in PBS and lysed using RIPA lysis buffer (Thermo Scientific #89900) with Halt^TM^ Protease Inhibitor Cocktail (Thermo Scientific #87786). For immunoblotting, the protein concentration was quantified utilizing Pierce^TM^ BCA Protein Assay Kit (Therma Fisher #23225), and 20 µg samples were prepared in 25 µl with 5x loading buffer. SEC fractions were prepared suspending 20 µl SEC sample with 5 µl of 5x loading buffer. Samples were resolved on 10% SDS-PAGE and transferred to PVDF membrane (BioRad #1620264). Membranes were incubated with ponceau, blocked (Li-Cor #927-5000), and incubated with primary antibodies (1:1,000) overnight at 4°C diluted in blocking buffer (Table S1). Membranes were rinsed, incubated with secondary antibody (1:20,000), and visualized on an iBright^TM^ FL 1500 Imaging System. Immunoblots were quantified utilizing ImageJ and normalized to the loading control.

### Mouse models and treatments

All mouse protocols were outlined in IACUC protocol #00003322 and were approved by the Oregon Health & Science University (OHSU) Department of Comparative Medicine. Subcutaneous tumors were established in 8-week-old female C57BL6 mice from Jackson Laboratories by injecting 1×10^6^ cells suspended 100 µL 1:1 cold PBS:Matrigel. Mice were euthanized at 30 days and tumors were excised and measured. Utilizing the established pancreatic orthotopic survival surgery, 4×10^4^ PDAC cells suspended in 20 µL 1:1 cold PBS:Matrigel were injected directly into the tail of the pancreas of 9-week-old male C57BL6 mice from Jackson Laboratories as the KPC-8069 cell line was derived from a male mouse. The peritoneum was closed sutured and the skin was closed using wound clips. After surgery, mice were injected with 0.1 mg/kg buprenorphine, provided wet food and monitored daily for a week for any signs of stress. Mice were weighed weekly and euthanized at between 7-21 days. For EV treatment, mice were injected intraperitoneally with 100 µL of EVs (1.5×10^9^ particles) suspended in PBS on alternating size every other day for the duration of the study. For functional assessment of mouse vasculature, DyLight^TM^ 649 I-B_4_ isolectin (Vector #DL-1208.5) was administered retro-orbitally 30 minutes prior to euthanasia. Mice were euthanized utilizing CO_2_ followed by cervical dislocation.

### Mouse tumor fixation, embedding, and sectioning

Mouse PDAC tumors were fixed in 4% PFA (Fisher #NC9288315) for 24 hours. Tumors for paraffin embedding were then rinsed with 70% ethanol and submitted to the OHSU Histopathology Core for paraffin embedding and 5 µm sectioning. Tumors for cryosectioning that were from mice perfused with lectin were moved from fixative to 30% sucrose for 24 hours. Tumors were then embedded in OCT and cryosectioned at 8 µm thickness and stored at −80°C until staining.

### Immunofluorescence staining of mouse tissues

Paraffin embedded mouse tissue sections were deparaffinized in xylene and rehydrated. Antigen unmasking was performed utilizing citric acid-based pH 6.0 solution (Vector Laboratories #H3300250) with high temperature for 20 minutes. For cryosections, tissues were rehydrated for 10 minutes in PBS. Tissues were then permeabilized, blocked for 1 hour, and incubated overnight with primary antibody (1:100) (Table S2) The following day tissues were rinsed, incubated with secondary antibody (1:500) for 1 hour, quenched for autofluorescence (Vector #SP-8400-15), stained with DAPI (1 µg/mL), and mounted (Invitrogen #P36934). Stained tissues solidified overnight at room temperature and were promptly scanned at 20x on a Zeiss Axio Scan.Z1 at the OHSU Advanced Light Microscopy Core. Images were quantified utilizing QuPath. Tumors were annotated and percent positive stained was calculated utilizing cell detection and object classifiers. Thresholding was determined based on secondary antibody only control sections.

### Tumor flow cytometry

Tumors were dissociated utilizing the GentleMACS, strained through a 40 µm filter, spun at 200 x g for 5 minutes, and then incubated with ACK lysis (Gibco™ #A1049201) for 1 minute. Cells were spun for 5 minutes at 200 x g and blocked in Mouse BD Fc Block™ (BD #553142) diluted 1:100 with fixable LIVE/DEAD™ (Invitrogen #L23105) diluted 1:500 in PBS for 30 minutes at 4°C. Following blocking, cells were rinsed and resuspended in staining solution with conjugated antibodies diluted in PBS with 2% FBS with 0.5 mM EDTA for 1 hour at 4°C in the dark. Cells were then rinsed twice, strained, resuspended in 200 µL with CountBright^TM^ beads (Invitrogen^TM^ #C36950), and run on a Cytek**^®^** Aurora 5-Laser Spectral Flow Cytometer. All experiments included single color controls and fluorescence-minus one control for gating. Flow cytometry data were analyzed utilizing FlowJo. Events were gated on cell size, singlets, live, followed by the gating strategy outlined in Supplemental Tables 3 and 4. Populations of interested were gated and cell numbers were calculated as percent of live, percent of CD45^+/-^, and cells per gram. Cells per gram was calculated by the following equation: 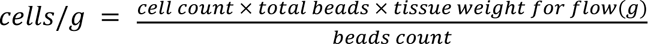. Fluorescence intensities were calculated by geometric mean.

### Extracellular vesicle RNA extraction

Pooled EV SEC fractions were lysed, and the RNA was extracted utilizing the qEV RNA Extraction Kit (Izon #RXT01). Per manufacturers instruction Lysis buffer A was heated at 60°C for 20 minutes prior to use. EV samples were lysed with 900 µL lysis buffer A and 125 µL lysis buffer B per 600 µL sample. RNA was isolated utilizing provided columns and eluted with 50 µL of elution buffer. Following isolation RNase inhibitor was added to each sample at a final concentration of 1 U/µL (Invitrogen #AM2682).

### Isobaric-labeling quantitative proteomics

Extracellular vesicles were pelleted via ultracentrifugation at 120,000 g for 2 hours. Samples were ultrasonicated, lysed in 5% SDS, 50 mM TEAB, and quantified before reducing, alkylating, and digesting 50 µg of each sample. 15 µg of peptide from each sample is dried, reconstituted in 25 µL of 100mM TEAB, pH8 and labeled using 6 channels (132N – 134C) of TMTPro16plex reagents (Thermo Scientific). The 6 labeled samples were pooled and analyzed on Orbitrap Fusion or Eclipse Tribrid mass spectrometer (Thermo Scientific) using synchronous precursor selection. The combined labeled peptide digests were separated using 20 online, high pH reverse phase fractions, followed by 140 minutes low pH reverse phase gradient. RAW instrument files were processed using COMET against a canonical human FASTA file, filtered for confident matches using the target/decoy method, and proteins inferred and grouped^60,61^. TMT reported ions were extracted and combined into protein abundance proxies using an established analysis pipeline. Jupyter notebooks with an R-kernel were used for quality control and edgeR statistical testing analyses^62^. The TMT experiment was normalized using the trimmed mean of M-values method followed by moderated test statistics and modeling. Protein comparisons required Benhamini-Hochberg correction *p*-values with an FDR < 0.1 for a protein to be considered differentially abundant.

### Reverse-phase protein array (RPPA)

Isolated EVs were shipped to Dr. Emanuel Petricoin for RPPA where EVs were printed in triplicate onto nitrocellulose-coated slides using a Quanterix 2470 Arrayer. Immunostaining was performed by probing each slide with a primary antibody targeting the protein of interest. Biotinylated goat anti-rabbit IgG (H + L) (1:7,500, Vector Laboratories) or rabbit anti-mouse IgG (1:10, DakoCytomation) were used as secondary antibodies. Signal amplification was carried out using a tyramide-based avidin/biotin system (DakoCytomation), followed by visualization with streptavidin-conjugated IRDye 680 (LI-COR). Negative controls were stained with secondary antibody only. Total protein was quantified using Sypro Ruby protein blot staining according to the manufacturer’s instructions (Molecular Probes). Total protein intensities for each sample were calculated by averaging the Sypro staining intensities of the three replicate spots.

### RNA-sequencing

RNA was isolated from cell pellets utilizing Quiagen RNeasy column purification. Isolated RNA from EVs and cells were quantified utilizing a NanoDrop spectrophotometer and RNA quality was assessed utilizing an Invitrogen Qubit RNA IQ assay. Isolated RNA was shipped to Novogene for poly-A enrichment, library construction, and RNA-sequencing. Post-sequencing analysis involved transcript quantification using Kallisto, which generated TPM and count matrices. Transcript-to-gene mapping was performed to aggregate transcript counts at the gene level using TxImport^63^. Differential expression analysis (DEA) was conducted using the R packages DESeq2 and edgeR to compare groups. For DESeq2, raw counts were normalized using variance-stabilizing transformation (VST) to ensure homoscedasticity across samples. The normalized data was analyzed using a generalized linear model (GLM) with subject and group as covariates to account for paired experimental designs. Genes with an adjusted *p*-value (FDR) < 0.05 were considered significantly differentially expressed. Results were annotated with Ensembl IDs and gene names using the biomaRt package. For edgeR, a similar GLM-based pipeline was implemented. Count data was normalized using the TMM (trimmed mean of M-values) method. Differential expression was assessed via likelihood ratio tests, and significant genes were identified based on FDR thresholds. Additional analyses included generating counts-per-million (CPM) matrices and exploratory data visualization.

### Single-cell RNA-sequencing analysis

Single-cell RNA sequencing (scRNA-seq) data from 115 PDAC samples were sourced from the Deeply Integrated Single-Cell Omics (DISCO) database^29^. A total 557,304 cells were utilized to investigate the relationship between *ELAVL1* gene expression in malignant cells and endothelial cell infiltration. Malignant cells were identified using the R package scATOMIC (v2.0.2) with the parameter “known_cancer_type = PAAD cell” and default settings for other parameters^64^. Spearman’s correlation analysis was conducted to assess the association between gene expression levels in tumor cells and cell-type infiltration in each sample.

### Extracellular vesicle import assays

EVs were labeled with the fluorescent lipid dye, PKH67. PKH67 was prepared per manufacturer’s instructions diluted 1:250 in Diluent C (Sigma, PKH67GL-1KT). EVs were incubated with the dye suspension for 10 minutes in the dark and then suspended over a cell pellet and immediately spun at 2000 x g for 5 minutes to remove excess dye from solution. PKH67 labeled EVs were suspended in media with 10% EV-depleted FBS and added to cells plated an 8 well chamber slides (Ibidi #80826). At the endpoint, media was aspirated, cells were rinsed with PBS and fixed utilizing 4% paraformaldehyde (PFA) for 15 minutes at room temperature. Cells were then rinsed, blocked for 30 minutes and stained with phalloidin (Biotium #00044, 1:100) and DAPI (1 µg/mL). Cells were imaged at 40x in 4×4 field of view tiles on a Zeiss Axio Observer. Images were analyzed on ImageJ by creating a mask around the phalloidin signal and measuring the intensity of the EVs within.

### Endothelial cell migration assay

HPaMECs or HUVECs were plated on 12-well cell culture inserts (Falcon #353181) that had been coated with fibronectin or Quick Coating Solution, respectively. PBS, PANC-1 WT, or PANC-1 HuR KO EVs were added to the well below the transwell and cells incubated for 6 hours. The top of the insert was then wiped clean and the cells that traveled to the lower membrane were fixed with 4% PFA, stained with 0.5% crystal violet, eluted with 5.8 M acetic acid and the absorbance was measured at 590 nm.

### Monolayer permeability assay

HPaMECs or HUVECs were plated in 12-well cell culture inserts (Falcon #353181) coated with fibronectin or Quick Coating Solution, respectively. Cells were incubated with media alone, PANC-1 HuR WT or KO EVs overnight with standard ECM media on the bottom of the transwell. The following day, FITC-dextran was added to the top chamber above the endothelial cell monolayer in addition to 100 µM H_2_O_2_ or EVs. After 6 hours, 100 µL media in triplicate per condition was taken from the bottom of the transwell, plated in a black 96 well plate, and the fluorescence was measured (λ_excitation_ = 498 nm, λ_emmission_ = 517 nm) to assess how much FITC-dextran passed through the monolayer.

### Tube formation assay

HPaMECs or HUVECs were treated for 24 hours with media alone, PANC-1 HuR WT or KO EVs. Treated cells were then plated in 15 well polymer coverslip slides (Ibidi 81506) coated with growth factor reduced Matrigel® (Corning #354230). Cells were incubated in a Keyence BZX Fluorescence Microscope with live imaging Tokai Hit Stage and imaged at 4x with phase contrast every 15 minutes for 24 hours. Images were analyzed using the WimTube image analysis platform.

### Ultrasound imaging

Tumor-bearing mice were anesthetized with isoflurane and the hair was removed. Mice were imaged on 3D color mode with MS250S transducer with 0.1mm step size on the Vevo 2100 Imaging System. 3D scans were quantified by drawing the tumor margin to determine percent vascularity on VevoLab.

### Transfection and lentiviral transduction

Cell lines were developed to express the PalmGRET EV reporter (pLenti-PalmGRET, Addgene #158221) utilizing lentivirus produced by transfected LentiX HEK293T cells. Following 24 hours of transduction, cells were selected with puromycin dihydrochloride (Sigma #P8833, 5 µg/mL).

### Statistics

All statistical analysis was performed utilizing GraphPad Prism (version 10.1.2). *In vitro* data are presented in bar plots with the representative replicate of 3 shown without single datapoints as mean ± standard deviation while *in vivo* data are presented with single datapoints as mean ± standard error of the mean of one of the two independent experiments. All experiments were performed in biological replicates, defined as independent experiments conducted on distinct biological samples, either separate cell culture experiments or individual animals. Exact *n* values for biological replicates are reported in the figure legends throughout. Statistical tests were unpaired two-tailed Student’s *t*-tests for comparing two groups and One-way ANOVA for multiple group comparisons.

## DATA AVAILABILITY

The raw bulk RNA-sequencing and processed data generated in this study have been deposited in the NCBI Gene Expression Omnibus database and are currently pending accession. The GEO accession number will be made available upon the acceptance of the manuscript for publication. The analyzed DESEQ data for RNA-sequencing are in Supplementary Data 1 (WT vs. HuR KO EV cargo) and 3 (HPaMEC treated with WT vs. HuR KO EVs). The raw tandem mass spectrometry data generated in this study have been deposited in the PRoteomics IDEntifications (PRIDE) Archive database and the accession will also be provided upon the acceptance of the manuscript for publication with dataset identifier PXD059674^65^. The analyzed proteomics data from WT vs. HuR KO EV cargo are in Supplementary Data 2. The remaining data supporting the results of this study are available from the corresponding author upon reasonable request. Scripts and resources used to generate the analyses are cited throughout the manuscript.

## AUTHOR CONTRIBUTIONS

Conceptualization: J.M.F., J.R.B.; software: M.R., K.H.; formal analysis: J.M.F, O.L., V.C., C.C.; investigation: J.M.F., Y.G., M.R., K.H., M.H., H.H., O.L., V.C.; writing – original draft: J.M.F., J.R.B.; writing – review & editing: J.M.F, Y.G., A.Q.B., M.H., R.C.S., K.T.B., J.R.B.; visualization: J.M.F.; supervision: Z.X., E.F.P., R.C.S., K.T.B., J.R.B.; funding acquisition: Z.X., E.F.P., R.C.S., K.T.B., J.R.B.

## DISCLOSURE OF INTEREST

The authors report no conflict of interest.

## ACKNOWLEDGEMENTS

We would like to thank all members of the Brody laboratory, the Brenden-Colson Center for Pancreatic Care, and the Knight Cancer Institute for their continued support and discussions regarding this work. This work was supported by the National Cancer Institute (NCI) of the National Institutes of Health (NIH) under awards R01 CA212600 (to J.R.B), U01 CA224012 (to R.C.S. & J.R.B), R21 CA263996 (to R.C.S. & J.R.B), and R01GM147365 (to Z.X.). American Association for Cancer Research (AACR) under award 15-90-25-BROD (to J.R.B), the Hirshberg Foundation, and the Brenden-Colson Center for Pancreatic Care. We thank and acknowledge the invaluable work performed by members of the OHSU Research Cores and Shared Resources. Electron microscopy was performed by Erin Stempinski at the Multiscale Microscopy Core, a member of the OHSU University Shared Resource Cores RRID:SCR_022652. Mass spectrometric analysis was performed by the OHSU Proteomics Shared Resource by Drs. Ashok Reddy and Phil Wilmarth with partial support from NIH core grants P30EY010572, P30CA069533, and S10RR02557. We thank all members of the Flow Cytometry Core, Advanced Light Microscopy Core, and Histopathology Shared Resource. We thank the OHSU Department of Comparative Medicine for their support in animal care, especially Allie Buckner for her hands on care of our animals.

## Notes

### Competing Interest Statement

The authors have declared no competing interest.

