## Supplemental Information for "Pancreatic cancer-intrinsic HuR regulates the pro-tumorigenic properties of extracellular vesicles"

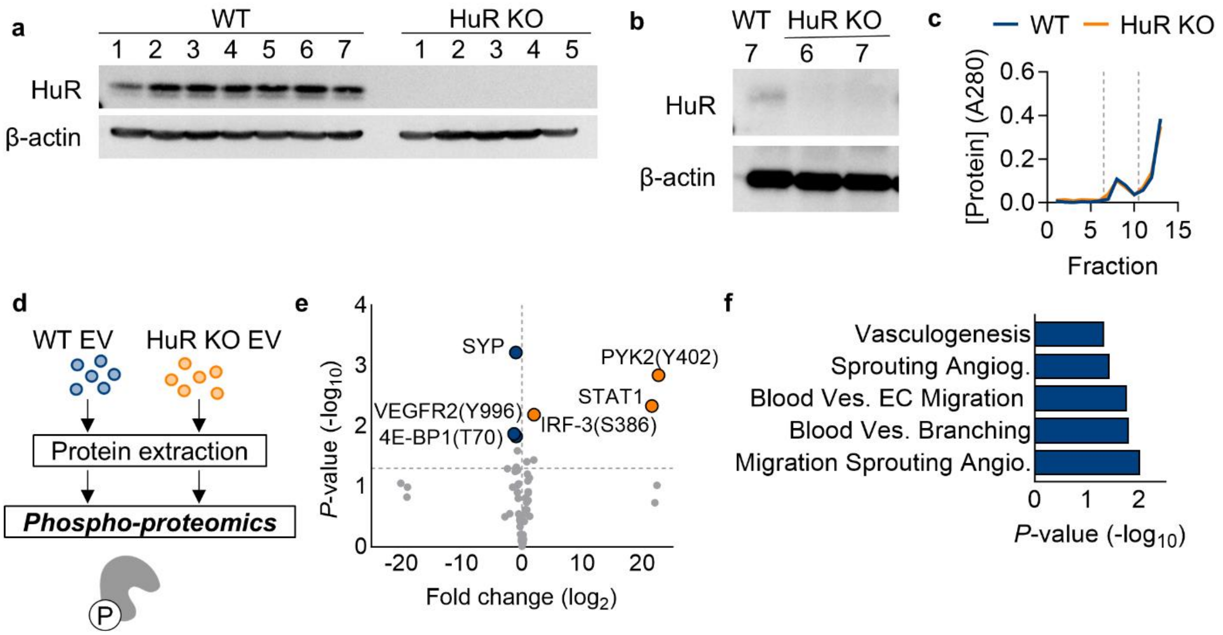


**Supplemental Figure 1: Validation of PANC-1 HuR KO and phospho-protein cargoes of PANC-1 EVs. (a)** Immunoblot validation of HuR expression in PANC-1 WT single cell clones 1-7 and HuR KO clones 1-5 probed for HuR and loading control β-actin. **(b)** Immunoblot validation of HuR expression in PANC-1 WT single cell clone 7 and HuR KO clones 6-7 probed for HuR and loading control β-actin. **(c)** Protein concentration of size exclusion chromatography fractions for PANC-1 WT (blue) vs. HuR KO (orange) EV isolations. **(d)** Schematic of phospho-proteomics on WT vs. HuR KO EVs. **(e)** Volcano plot of differentially abundant proteins and phospho-proteins (amino acid phosphorylated listed) enriched in WT (left, blue) vs. HuR KO (right, orange) EVs. **(f)** Gene ontology analysis of proteins and phospho-proteins enriched in WT EVs.

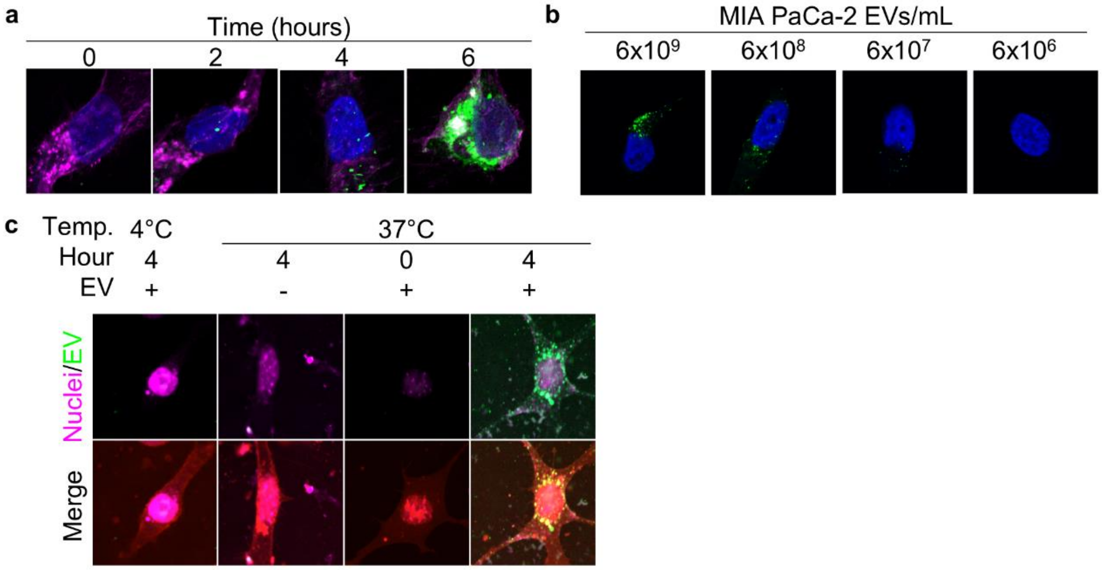


**Supplemental Figure 2: Validation of EV import imaging *in vitro*. (a)** MIA PaCa-2 cells treated with PKH67 labeled MIA PaCa-2 EVs (green) and collected at 0-6 hours and stained for nuclei (DAPI, blue) and the cell surface (wheat germ agglutinin, magenta). **(b)** MIA PaCa-2 cells treated with a titration of $6x10^6$ – $6x10^9$ PKH67 labeled MIA PaCa-2 EVs (green) and collected at 4 hours and stained for nuclei (DAPI, blue). **(c)** Cancer associated fibroblasts treated with PKH67 labeled MIA PaCa-2 EVs (green) for 0 and 4 hours at 37°C and 4°C or PKH67 alone (EV-) for 4 hours.

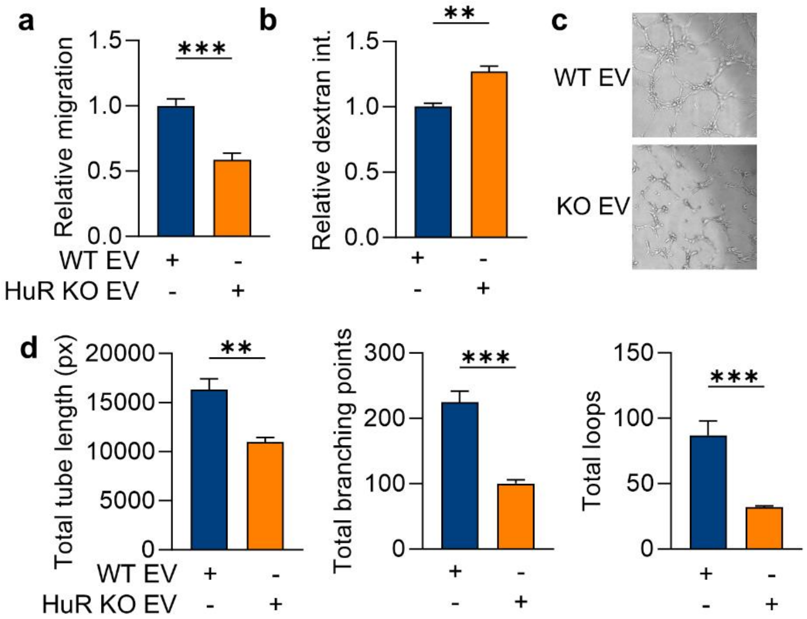


**Supplemental Figure 3: Repeating *in vitro* EV treatments with an additional endothelial cell line. (a)** Transwell migration of HUVECs when treated with PANC-1 WT vs. HuR KO EVs over 24 hours and quantified with crystal violet staining ($n$ = 3). Functional analysis of HUVECs treated with media alone, PANC-1 WT or HuR KO EVs for 24 hours and monitored for **(b)** monolayer permeability quantified by dextran movement across the endothelial cell monolayer and **(c-d)** tube formation quantified for total tube length (px), total branching points and total loops ($n$ = 3). $P$ values were calculated using an unpaired two-tailed Student's $t$-test. *, $P < 0.05$; **, $P < 0.01$; ***, $P < 0.001$; ns, not significant.

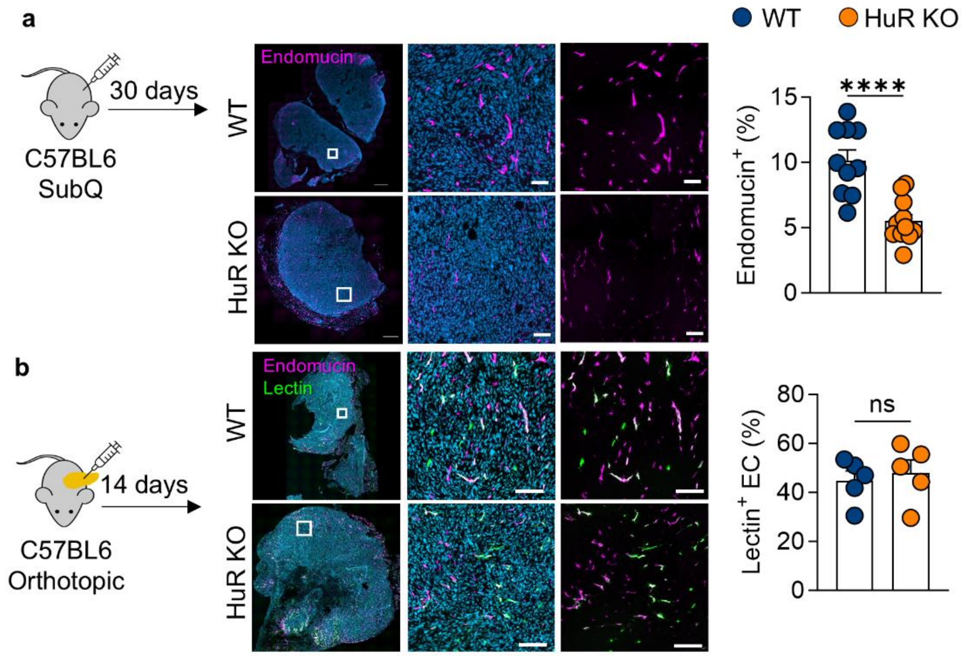


**Supplemental Figure 4: Additional mouse model staining. (a)** KPC HuR WT vs. KO cells subcutaneously implanted into the flank of C57BL6 mice and euthanized after 30 days and stained for endomucin (magenta) and nuclei (DAPI, blue) (*n* = 7). **(b)** Orthotopic WT vs. HuR KO tumors perfused with lectin (green) and stained for nuclei (DAPI, blue) and endothelial cells (endomucin, magenta) (*n* = 5). Scale bars = 100 µm. *P* values were calculated using an unpaired two-tailed Student's *t*-test. *, *P* < 0.05; **, *P* < 0.01; ***, *P* < 0.001; ns, not significant.

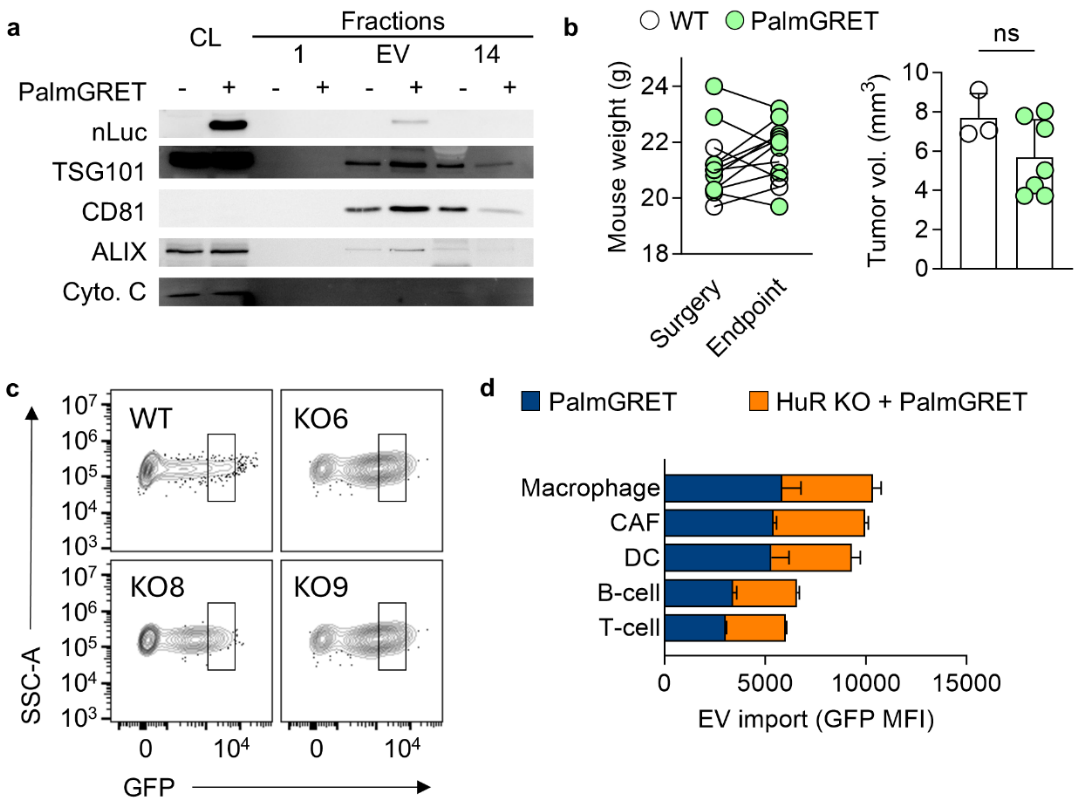


**Supplemental Figure 5: Validation of PalmGRET reporter in KPC cells. (a)** Immunoblot of cell lysates (CL) and SEC fractions 1, EV and 14 from KPC WT and PalmGRET cells and probed for the PalmGRET reporter expression (nLuc), classical EV markers (TSG101, CD81, ALIX) and negative control cytochrome C. **(b)** Mouse bodyweight and tumor volume from KPC WT ($n$ = 3) and PalmGRET ($n$ = 7) orthotopic tumors. **(c)** Flow contour plots of transduced KPC cells with PalmGRET with gates for fluorescence-activated cell sorting based on GFP intensity. **(d)** EV import across all stromal cell in flow cytometry panel in PalmGRET vs. PalmGRET HuR KO tumors as reported by GFP geometric mean (MFI). $P$ values were calculated using an unpaired two-tailed Student's $t$-test. *, $P < 0.05$; **, $P < 0.01$; ***, $P < 0.001$; ns, not significant.

| **Supplemental Table 1: Immunoblotting antibody information** | | |
|---|---|---|
| **Marker** | **Source** | **Catalog #** |
| HuR | Santa Cruz | sc-5261 |
| Vinculin | Santa Cruz | sc-73614 |
| TSG101 | Abcam | ab125011 |
| CD81 | Santa Cruz | sc-166029 |
| ALIX | Cell Signaling | 2171S |
| Cytochrome C | Cell Signaling | 11940S |
| β-actin | Cell Signaling | 4967S |
| nLuc | Promega | N7000 |

| **Supplemental Table 2: Immunofluorescence antibody information** | | |
|---|---|---|
| **Marker** | **Source** | **Catalog #** |
| Ki67 | Cell Signaling | 12202S |
| Endomucin | eBiosciences | 14-5851-82 |
| DLL4 | Novus Biologicals | AF1389-SP |
| ICAM-1 | Thermo Scientific | 14-0541-82 |

| **Supplemental Table 3: Flow cytometry antibody information** | | | | |
|---|---|---|---|---|
| **Marker** | **Fluorophore** | **Dilution** | **Source** | **Catalog #** |
| MHC-II | AF700 | 1:100 | BioLegend | 107622 |
| ICAM-1 | APC-Fire750 | 1:100 | BioLegend | 116126 |
| CD19 | BUV805 | 1:200 | BD | 568287 |
| CD90.2 | BUV395 | 1:100 | BD | 565257 |
| Live/Dead | LD Blue | 1:500 | Invitrogen | L23105 |
| F4/80 | BV421 | 1:100 | BioLegend | 123137 |
| CD11c | BV650 | 1:100 | BioLegend | 117339 |
| CD31 | BV785 | 1:100 | BioLegend | 102435 |
| PDPN | PE | 1:100 | BioLegend | 127408 |
| CD45.2 | PE Cy7 | 1:100 | BioLegend | 109830 |

| **Supplemental Table 4: Flow cytometry gating strategy for orthotopic pancreatic tumors** | |
|---|---|
| **Cell Type** | **Gating (of live single cells)** |
| EC | $CD45^{-}CD31^{+}$ |
| CAF | $CD45^{-}CD31^{-}PDPN^{+}$ |
| T cell | $CD45^{+}MHCII^{-}CD90^{+}$ |
| B cell | $CD45^{+}MHCII^{+}CD19^{+}$ |
| DC | $CD45^{+}MHCII^{+}CD11c^{+}$ |
| Macrophage | $CD45^{+}MHCII^{+}F4/80^{+}$ |
